# Evaluation of TRPM8 and TRPA1 participation in dental pulp sensitivity to cold stimulation

**DOI:** 10.1101/212688

**Authors:** Benoit Michot, Caroline Lee, Eugene Podborits, Jennifer L. Gibbs

## Abstract

Sensory neurons innervating the dental pulp have unique morphological and functional characteristics compared to neurons innervating other tissues. Stimulation of dental pulp afferents whatever the modality or intensity of the stimulus, even light mechanical stimulation that would not activate nociceptors in other tissues, produces an intense pain. These specific sensory characteristics could involve receptors of the Transient Receptor Potential channels (TRP) family. In this study, we evaluated 1) the expression of TRPA1 and TRPM8 receptors in trigeminal ganglion neurons innervating the dental pulp compared to sensory neurons innervating the oral mucosa or the skin of the face, and 2) the involvement of these receptors in dental pulp sensitivity to cold stimulation. We showed a similar expression of TRPM8 and CGRP in sensory neurons innervating the dental pulp, the skin or the buccal mucosa. On the contrary, TRPA1 was expressed in a higher proportion of neurons innervating the mucosa (43%) than in neurons innervating the dental pulp (19%) or the skin (24%). Moreover, neurons innervating the dental pulp had a higher proportion of large neurons (24%) compared to neurons innervating the skin (8%) or the mucosa (10%). The evaluation of trigeminal ganglion neuron sensitivity to TRPM8 agonist, TRPA1 agonist and cold stimulation, showed that a significant proportion of neurons innervating the skin (10%) or the mucosa (37%) were sensitive to cold stimulation but insensitive to TRPM8 and TRPA1 activation. Similarly, the application of a cold stimulation on the tooth induced an overexpression of cFos in the trigeminal nucleus that was not prevented by administration of a TRPA1 antagonist or the genetic deletion of TRPM8. However, the pretreatment with the local anesthetic carbocaine abolish the cold-induced cFos overexpression. In conclusion, the unique sensory characteristics of the dental pulp would be independent to TRPA1 and TRPM8 expression and functionality.

## 1. Introduction

Dental pulp is a highly innervated tissue in which sensory afferents have unique characteristics compared to other tissue. In the pulp, although most peripheral nerve endings appear to be small unmyelinated C and small myelinated A delta nociceptors, they originate from large diameter myelinated fibers in the trigeminal ganglion (Cadden et al. 1983; Paik et al. 2009; Fried et al., 2011; Gibbs et al. 2011). These sensory afferents have characteristics of low threshold mechanosensor, they express NF200, parvalbumin and upregulate NPY after injury (Fried et al., 1989; Itotagawa et al.,1993; Ichikawa et al. 1995), but they also express for a part of them markers of nociceptors such as TrkA, CGRP and TRPV1 (Yang et al. 2006; Gibbs et al. 2011). Further, stimulation of dental pulp afferents whatever the modality or intensity of the stimulus, even light mechanical stimulation that would not activate nociceptors in other tissues, produces an intense pain (Dababneh et al. 1999).

In general, detection of thermal and mechanical stimuli are attributable to receptors of the Transient Receptor Potential (TRP) channels family (Julius 2013). Among these receptors, TRPA1 and TRPM8 are important in the activation of sensory neurons by cold temperatures (McKemy et al. 2002; Peier et al. 2002; Story et al. 2003). TRPA1 is a polymodal receptor activated by noxious cold stimulation (<17°C), chemical irritants (acyl isothiocyanate, acrolein, formaline…) and appears to be involved in mechanosensation as well (Story et al. 2003; Bautista et al. 2006; Kwan et al. 2006; McNamara et al. 2007). Its role in cold perception is not fully understood but pharmacological blockade and genetic deletion of TRPA1 reduces cold sensitivity in physiological, inflammatory and neuropathic conditions (Kwan et al. 2006; Karashima et l. 2009; Chen et al. 2011; Nassini et al. 2011). In contrast, TRPM8 is expressed in a distinct neuronal subpopulation and has is activated at non-noxious cold temperature (<25°C; Peier et al. 2002; Story et al. 2003). While TRPM8 is clearly involved in cool temperature perception, it may also mediate noxious cold sensation, as TRPM8 antagonists reduce cold hypersensitivity in neuropathic pain models (Knowlton et al. 2013; Patel et al. 2014).

Although both TRPA1 and TRPM8 are expressed in sensory neurons innervating the dental pulp (Park et al. 2006), their involvement in dental pulp sensitivity to cold and whether they contribute to the sensory characteristics of the dental pulp is, to date, not known. In this study, we examined the expression of TRPM8 and TRPA1 in sensory neurons innervating the dental pulp compared to neurons innervating facial skin or the oral mucosa. We also evaluated the participation of TRPA1 and TRPM8 in dental pulp detection of noxious cold stimulation.

## 2 Materials and Methods

### 2.1 Animals

All experiments were approved by the Institutional Animal Care and Use Committee at the New York University College of Dentistry and followed the guidelines provided by the National Institutes of Health Guide for the Care and Use of Laboratory Animals. Experiments were performed on 10-12 weeks old male and female C57Bl6 and TRPM8 knock-out (*Trpm8*^*tm1Apat*^-GFP-tagged) mice purchased from Jackson Laboratory. Animals were housed 2-5/cage in an environment with controlled 12h/12h light/dark cycles and had free access to food and water.

### 2.2 Retrolabeling

Mice were anaesthetized with 2% isoflurane and injected with 2μl of Fluorogold (2.5% in saline) or 1,1’-Dioctadecyl-3,3,3’,3’-Tetramethylindocarbocyanine Perchlorate (DiI, 2.5% in ethanol) either intradermally in the skin of the cheek or submucossally in the buccal mucosa to label the respective innervating neurons. Injections were repeated 2-3 times in each region. Retrolabeling of dental pulp innervating neurons was performed in mice deeply anaesthetized by an intraperitoneal injection of ketamine-xylazine (100-10mg/kg). The enamel of the first molar in each side of the jaw was removed with a ¼ round dental bur at low speed to expose dentinal tubules without exposing the dental pulp to the oral environment. The tooth surface was washed, and the smear layer was removed with 35% phosphoric acid gel. Fluorogold or DiI (1-2μL) was applied three times with a 5-minute drying interval between each application. At the end of each procedure mice were treated once a day for three days with carprophen (5mg/kg, i.p.) to avoid the development of a local inflammation that could affect protein expression in sensory neurons.

### 2.3 TRPA1 and CGRP immunostaining

Seven days after retrolabeling sensory neurons with Fluorogold, homozygotes and hererozygotes *Trpm8*^*tm1Apat*^-GFP-tagged and wild type mice were euthanized and perfused intracardially with 10ml of phosphate-buffered saline (PBS 1X) followed by 30ml of 10% formaline in PBS. The trigeminal ganglia were removed, post-fixed for 30min in 10% formaline and cryoprotected overnight in PBS with 30% sucrose. Trigeminal ganglia were cut (16μm thick sections) with a Cryostat (-20°C) and collected on superfrost glass slides. Sections were washed for 30min in PBS, followed by 1h incubation in a blocking solution containing 0.3% Triton X100 and 5% normal goat serum in PBS. Sections were then incubated at 4°C for either 3 days with rabbit anti-TRPA1 antibody (1:10000; AB58844; Abcam) or 1 day with chicken anti-GFP antibody (1:10000; GFP-1020; Aves Labs). After a 30-min wash in the blocking solution, samples were incubated for 2h with the secondary antibody Alexa Fluor 488-conjugated goat anti-rabbit (1:700; Life Technology) or Alexa Fluor 488-conjugated goat anti-chicken (1:700; Life Technology). Then, sections were washed in blocking solution for 30min and incubated at 4°C overnight with mouse anti-CGRP antibody (1:8000; C7113, Sigma). The following day, sections were washed for 30min in PBS and successively incubated for 1h in the blocking solution and 2h with the secondary antibody Alexa Fluor 546-conjugated goat anti-mouse (1:700; Life Technology). After a final 30min wash, sections were coverslipped. Photomicrographs of trigeminal ganglia were collected with a fluorescent inverted microscope (Nikon Eclipse Ti). The proportion of Fluorogold retrolabeled neurons that expressed TRPA1, GFP and/or CGRP was quantified using the NIH Image J analysis Software. The cell body of all fluorogold retrolabeled neurons was delineated, then, after background substraction, neurons with more than 50% of the cell body area stained for TRPA1, GFP and/or CGRP were considered immunoreactive for these proteins and quantified.

### 2.4 Cold stimulation and cFos immunostaining

Female C57Bl6 mice pretreated with either vehicle, the TRPA1 antagonist HC030031 (dissolved in DMSO, 100mg/kg, i.p.) or carbocaine (3% in saline, 10μl injected periodontally) and TRPM8 knockout mice or wild type mice were anaesthetized with ketamine-xylazine. The jaw was opened and a unilateral cold stimulation was applied on the first molar using a small piece of cotton cooled with Frigi-dent (Ellman International, Inc). The stimulation was repeated 15 times over a 30-min period. Three hours after the beginning the stimulation mice were perfused intracardially with formaline, then the brainstem was collected, post-fixed for 3h and cryoprotected as described in part 2.6. The brainstem was cut (40μm thick sections) in a Cryostat (-20°C) and collected in containers filled with PBS. Every 5^th^ section was selected for staining. Floating sections were incubated 1h in a blocking solution containing 0.3% Triton X100 and 5% normal goat serum in PBS and then incubated overnight with rabbit anti-cFos antibody (1:30000, PC38, Calbiochem). After 30min wash with the blocking solution, sections were incubated for 1h with the secondary antibody biotinylated goat anti-rabbit (1:700; Vector). After a 30-min wash with PBS, the sections were successively incubated for 30min in 0.3% H_2_O_2_ in PBS and for 1h with the avidin-biotinylated-horseradish-peroxidase (ABC Vectastain kit Elite, Vector). Then they were washed 10min with PBS and stained black/gray with 3,3′-Diaminobenzidine (DAB kit, Vector). After a final 30min wash in PBS, sections were mounted on superfrost glass slides and coverslipped. The number of c-Fos immunoreactive neurons was quantified in both ipsilateral and contralateral trigeminal nucleus, in sections form caudalis through interpolaris region. The different regions of the trigeminal nucleus were identified according to the Allen Brain Atlas.

### 2.5 Trigeminal ganglion neuron culture

Trigeminal ganglia from DiI retrolabeled C57Bl6 mice were collected in F12 culture media and cut in ten pieces each. Then, they were incubated for 20min with papain (40units/ml) in Hank’s Balanced Salt Solution (HBSS) without calcium. After 2min centrifugation at 180g the supernatant was removed, and trigeminal ganglia incubated for 20min in collagenase-dispase solution (3.33-4.66mg/ml in HBSS without calcium). After 4min centrifugation at 400g the supernatant was removed. Neurons were resuspended in F12 culture media and centrifuged for 6min at 460g to remove the excess of collagenase-dispase solution. Then, neurons were suspended in F12 media supplemented with 5% Fetal Bovine Serum, mechanically dissociated with a P200 pipet and seeded into poly-D-Lysine-coated 25mm round glass coverslips. After 3h incubation at 37°C in a humidified atmosphere containing 5% CO_2_, neuronal culture was processed for calcium imaging experiment.

### 2.6 Single-cell calcium imaging

Trigeminal ganglion neurons cultured on round glass coverslips were washed for 10min in HBSS and loaded in the dark with the fluorescent calcium indicator Fura-2-acetoxy-methyl ester (Fura2, 5μM) for 45min at room temperature. Then, coverslips were washed for 10min in HBSS and mounted in a microscope chamber. Loaded cells were excited successively (2Hz) for Fura2 at 340 and 380nm for 200ms and emitted fluorescence was monitored at 510nm using a charged coupled device sensor camera coupled to an inverted Nikon Eclipse Ti microscope. Fluorescence intensities from single cells excited at the two wavelengths were recorded separately, corrected for the background and the fluorescence ratio (F340/F380) was calculated using the software NIS Elements-AR version 4.0. All neurons were stimulated for 1min successively with the TRPA1 agonist acyl-isothiocyanate (AITC, 250μM), the TRPM8 agonist menthol (250μM) and ice cooled HBSS (0-2°C). At the end of each treatment, the solution was removed, and neurons were washed for 3min with HBSS before starting the following treatment. At the end of each experiment KCl (75mM) was applied to identify healthy neurons. Only neurons retrolabeled with DiI that responded to KCl were used for the analysis. A positive neuronal response was defined as a 20% increase of the ratio F340/F380 relative to the pretreatment F340/F380 value.

### 2.7 Statistical analyses

Statistical analyses were performed with GraphPad Prism 5 software. Immunostaining of TRPM8, TRPA1 and CGRP, and calcium imaging data are expressed as percentage of immunoreactive or responsive neurons relative to the number of Fluorogold/DiI positive neurons and were analysed with Chi-square test. Results of cFos immunostaining are expressed as a mean+S.E.M of the number immunoreactive neurons/side/section/animal and were analyzed with a 2-way ANOVA, followed by Bonferroni post hoc test. The significance level was set at p<0.05.

## 3 Results

### 3.1 Expression of TRPM8 and TRPA1 in sensory neurons innervating the facial skin, the buccal mucosa or the dental pulp

The dental pulp has unique sensory characteristics that could be dependent on TRP channel expression in pulp innervating afferents. We addressed this question comparing the expression of the cold sensitive receptors TRPM8 and TRPA1 in sensory neurons innervating the dental pulp, the skin of the cheek or the buccal mucosa (Fig.1, 2). TRPM8 receptors were expressed in 5.7% of neurons innervating the dental pulp which was similar to the frequency of TRPM8 expression in neurons innervating the facial skin and the buccal mucosa (3.3% and 6.7% of TRPM8 immunoreactive neurons respectively, Fig.3A). Similarly, CGRP, a marker of small peptidergic nociceptors, was expressed in the same proportion of labeled sensory neurons whatever the innervating tissue (skin: 17.8%, mucosa: 14.3%, and dental pulp: 14.9%; Fig.3C). Interestingly, TRPA1 was expressed in a higher proportion of neurons innervating the mucosa than neurons innervating the dental pulp (43.0% versus 18.9%, p=0.0008, Chi-square test; Fig.3B) or the skin (43.0% versus 24.6%, p<0.0001, Chi-square test; Fig.3B). Moreover, we showed that a lower, but not statistically significant, proportion of neurons innervating the dental pulp co-expressed CGRP and TRPA1 compared to neurons innervating the skin (3.7% versus 13.7%, p=0.091, Chi-square test; Fig.3D) or the mucosa (3.7% versus 12.7%, p=0.15, Chi-square test, Fig.3D). To further characterized neurons innervating the dental pulp we evaluated the size distribution of neurons (Fluorogold positive) innervating the dental pulp, the skin or the mucosa, and neurons expressing TRPA1 and CGRP that innervate each tissue. The proportion of large neurons (>1200μm2) is significantly higher in the neuronal population innervating the dental pulp (24.1% of large neurons) than in the neuronal population innervating the skin (8.7% of large neurons, p<0.0001, Chi-square test; Fig.3E) or the mucosa (10.7% of large neurons, p=0.0002, Chi-square test, Fig.3E). The comparison of the size distribution of CGRP or TRPA1 immunoreactive neurons showed no difference between neurons innervating the dental pulp, the skin or the mucosa (Fig.3F, 3G).

**Figure 1:**
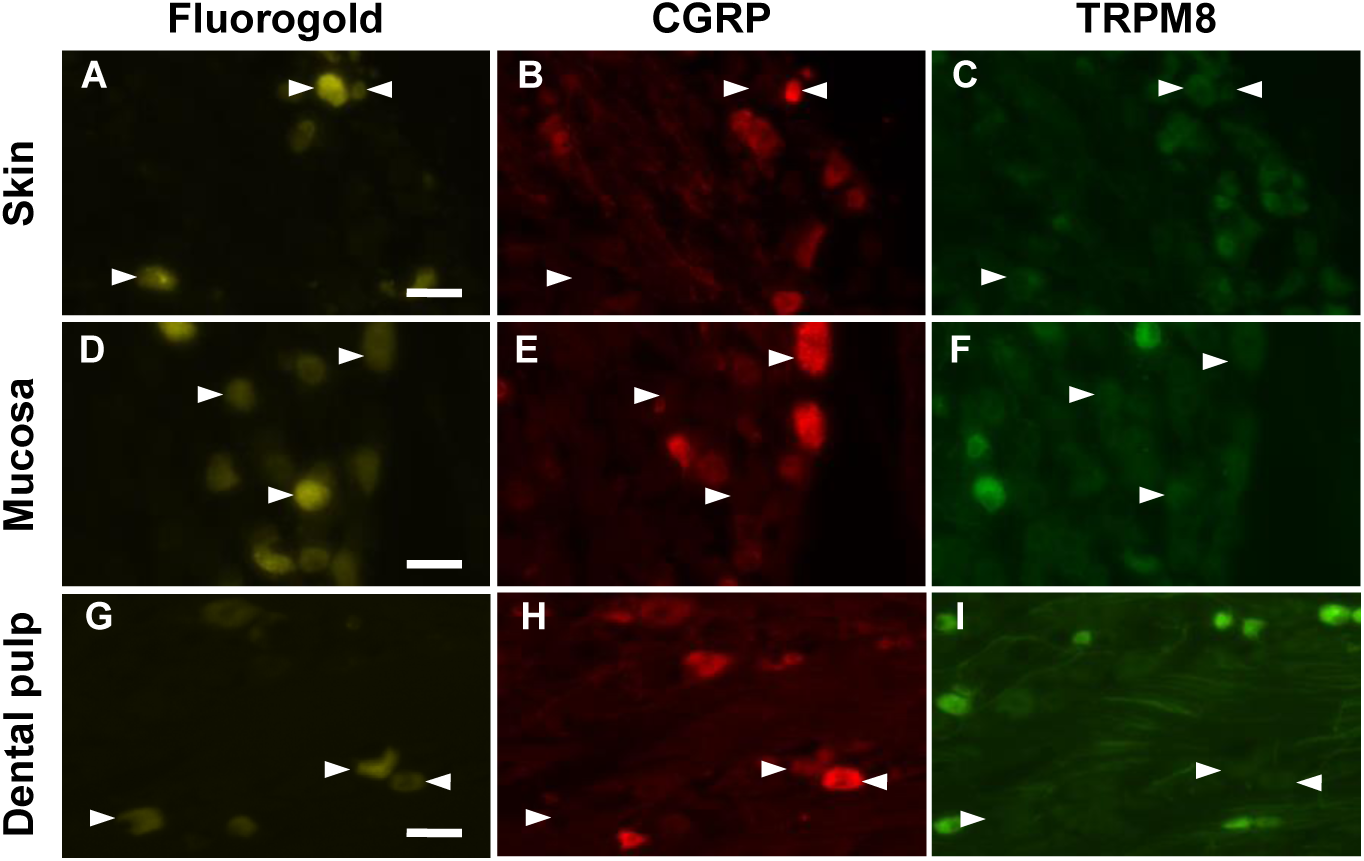
Expression of TRPM8 and CGRP in neurons innervating the facial skin, the oral mucosa or the dental pulp. Trigeminal sensory neurons innervating either the facial skin (A, B, C), the oral mucosa (D, E, F) or the dental pulp (G, H, I) were retrolabeled with Fluorogold (yellow) in *Trpm8*^*tm1Apat*^-GFP-tagged mice. Seven days later trigeminal ganglia were collected and immunostained for CGRP (red). Neurons expressing TRPM8 are stained green. Arrowheads indicate fluorogold retrolabeled neurons. Scale bar: 50μm.

**Figure 2:**
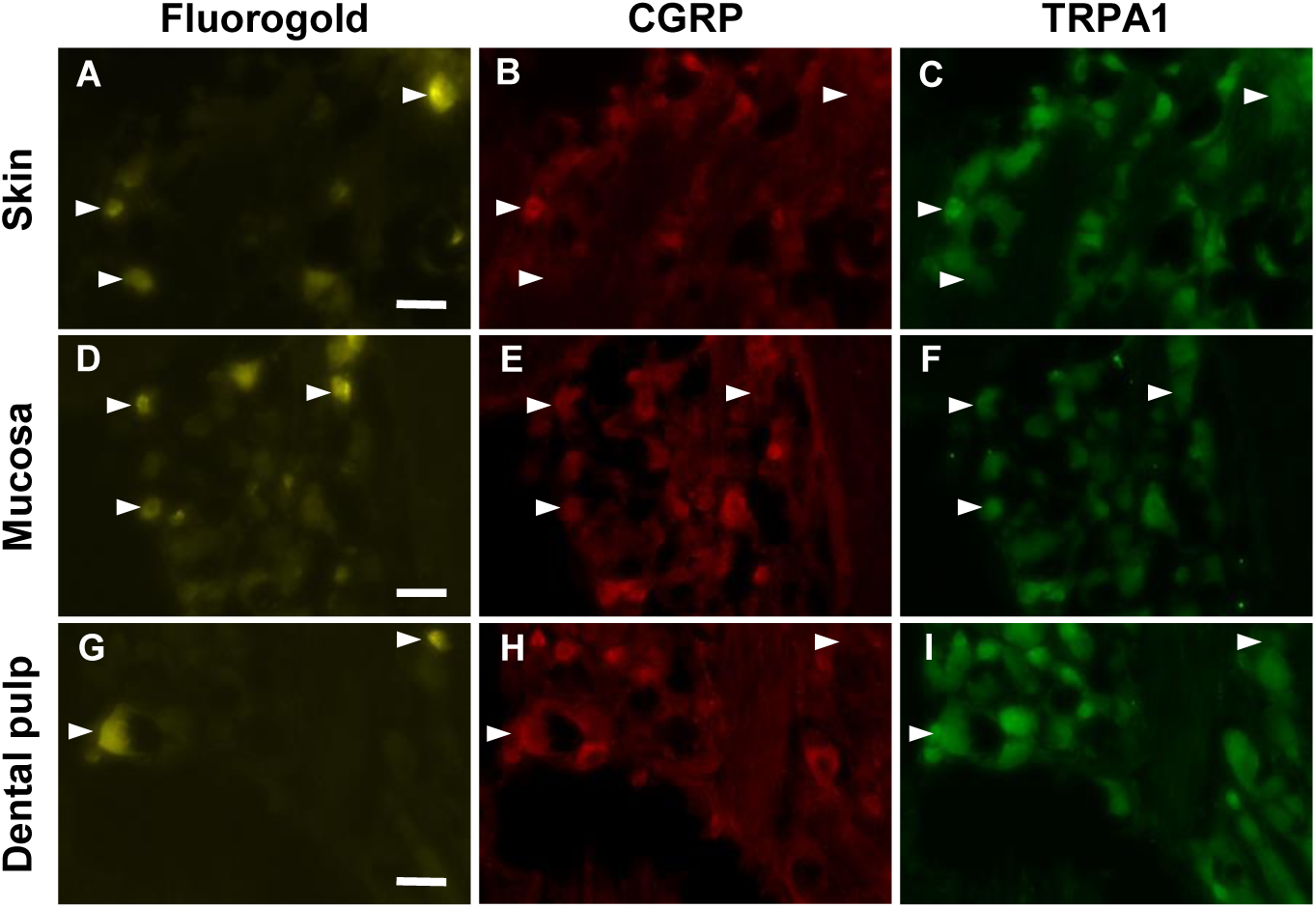
Expression of TRPA1 and CGRP in neurons innervating the facial skin, the oral mucosa or the dental pulp. Trigeminal sensory neurons innervating either the facial skin (A, B, C), the oral mucosa (D, E, F) or the dental pulp (G, H, I) were retrolabeled with Fluorogold (yellow). Seven days later trigeminal ganglia were collected and immunostained for CGRP (red) and TRPA1 (green). Arrowheads indicate fluorogold retrolabeled neurons. Scale bar: 100μm

**Figure 3:**
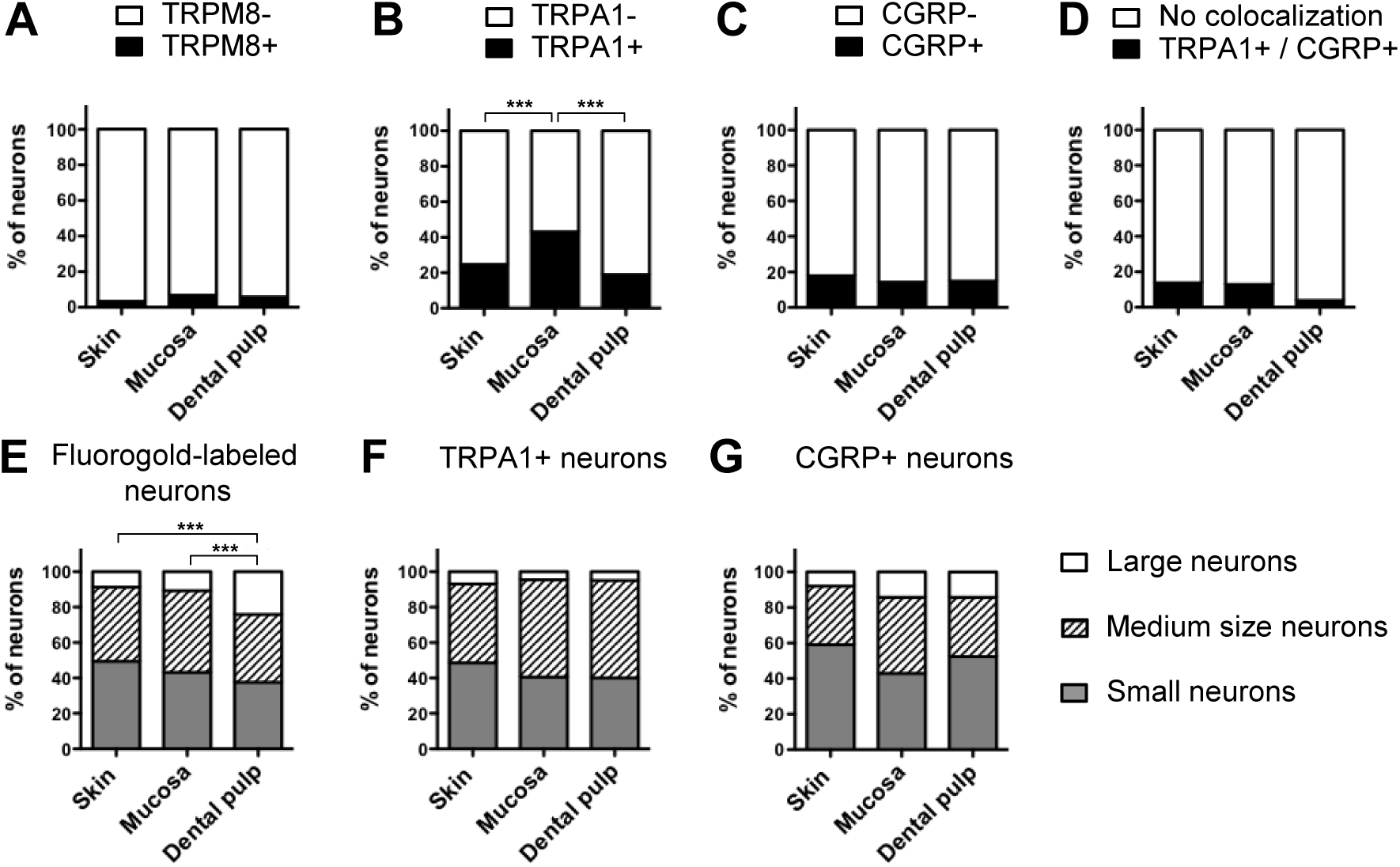
Quantitative analysis of TRPM8, TRPA1 and CGRP expression in trigeminal sensory neurons innervating the facial skin, the oral mucosa or the dental pulp. Proportion of Fluorogold retrolabeled trigeminal neurons that express, A) TRPM8, B) TRPA1, C) CGRP, and D) both TRPA1 and CGRP. E), proportion of small, medium and large neurons in the neuronal population (retrolabeled with Fluorogold) innervating either the skin, the mucosa or the dental pulp. F) and G), proportion of small (<600μm^2^), medium (600-1200μm^2^) and large neurons (>1200μm^2^) expressing CGRP or TRPA1 in the neuronal population innervating either the skin, the mucosa or the dental pulp. *** p<0.001, Chi-square test, n=106-410 neurons.

### 3.2 TRPM8 and TRPA1 receptors are independent to dental pulp sensitivity to cold stimulation

The application of a unilateral cold stimulation on the first molar induced neuronal activation detected through the ipsilateral increase of cFos expression in the trigeminal nucleus (Fig.4A, 4B). In the caudalis and the interpolaris part of the trigeminal nucleus the number of cFos imunoreactive neurons was not different in the ipsilateral versus contralateral side. However, in the transition zone between the caudalis and the interpolaris regions, the number of cFos immunoreactive neurons was significantly higher in the ipsilateral compared to the contralateral side (34.8+/-5.1 and 10.5+/-3.0 respectively; 2-way ANOVA; side: p=0.0018, F=10.6, DF=1; region: p=<0.0001, F=16.28, DF=2; interaction: p=0.0002, F=9.59, DF=2; Bonferroni’s test, transition zone ipsi versus contra p<0.001; Fig.5C).

**Figure 4:**
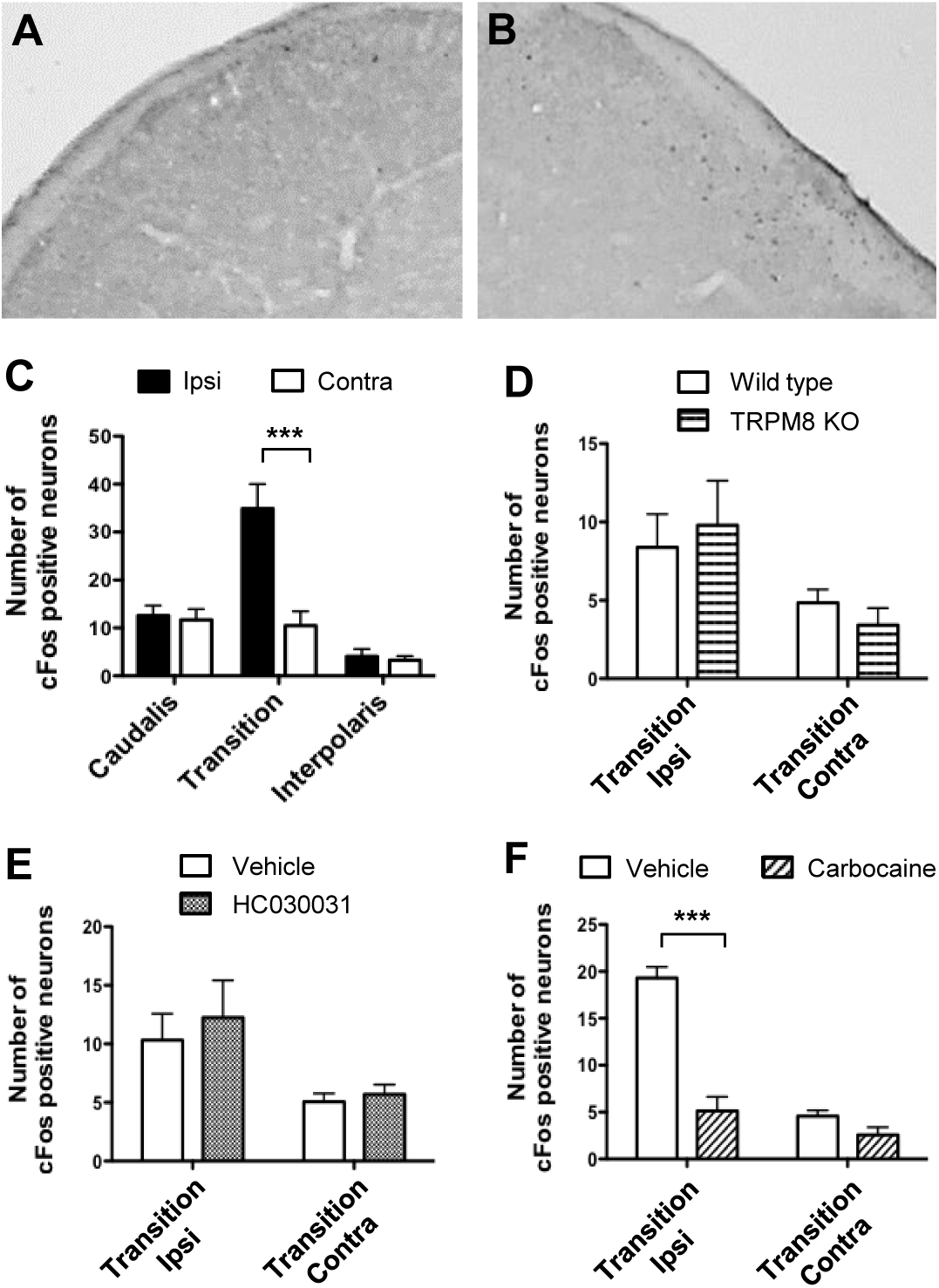
Modulation of cold-induced cFos upregulation in the transition zone of the trigeminal nucleus. Representative photo-micrographs of cFos immunoreactive neurons in A) contralateral and B) ipsilateral transition zone of the trigeminal nucleus of mice which underwent unilateral cold stimulation of the 1^st^ molar. C), quantitative analysis of cFos expression in the caudal part, the transition zone and the interpolaris part of the trigeminal nucleus. Effects of genetic deletion of TRPM8 (D), TRPA1 antagonist (E) and Cabocaine (F) on cold-induced upregulation of cFos in the trigeminal nucleus transition zone. Data are expressed as mean + S.E.M. of the number of neurons/section/animal. *** p<0.001, Bonferroni’s test, n=5-13/group.

**Figure 5:**
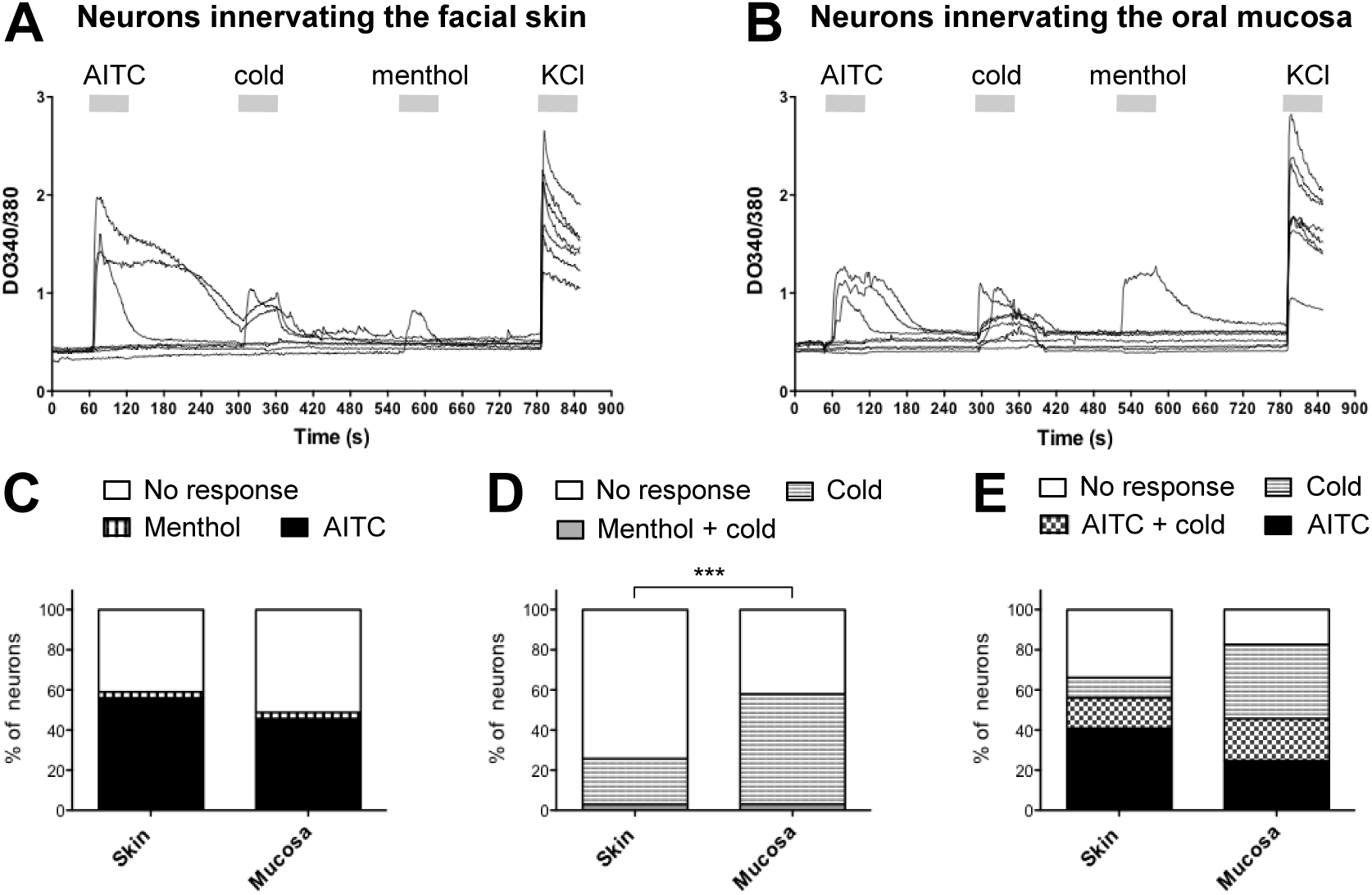
Effects of AITC, cold stimulation and menthol on sensory neurons innervating the facial skin or the oral mucosa. A) and B), each trace represents intracellular calcium levels in one neuron (retrolabeled with DiI) innervating the facial skin or the oral mucosa that was treated with AITC (250μM), cold HBSS buffer, menthol (250μM), and KCl (75mM). Horizontal grey bars indicate the duration of each treatment. Bar-graphs show the proportion of neurons responding to, C) AITC or menthol, D) menthol or cold HBSS buffer, E) AITC and/or cold HBSS buffer. Note that no neurons were responsive to both AITC and menthol. *** p<0.001, Chi-square test, n=66-162 neurons.

The involvement of TRPM8 and TRPA1 receptors in dental pulp sensitivity to cold stimulation was evaluated using TRPM8 KO mice and the TRPA1 antagonist, HC030031 (100mg/kg, i.p.). In TRPM8 KO mice, cold stimulation increased the number of cFos immunoreactive neurons in the ipsilateral trigeminal nucleus transition zone (ipsi: 9.8+/-3.4 versus contra: 3.4+/-1.1) but the increase was not different to the number of cFos immunoreactive neurons in wild type mice (TRPM8 KO ipsi: 9.8+/-3.4 neurons versus WT ipsi: 8.4+/-2.1 neurons; 2-way ANOVA; side: p=0.014, F=7.26, DF=1; genotype: p=0.99, F=0.00001, DF=1; interaction: p=0.45, F=0.59, DF=1; Fig.4D). Similarly, HC030031 did not prevent cold-induced upregulation of cFos in the ipsilateral compared to the contralateral trigeminal nucleus (2-way ANOVA; side: p=0.008, F=7.99, DF=1; treatment: p=0.54, F=0.37, DF=1; interaction: p=0.76, F=0.096, DF=1; Fig.4D). On the contrary, the administration of carbocaine (3% in saline) adjacent to the stimulated tooth reduced the cold-induced increase of the number of cFos immunoreactive neurons in the trigeminal nucleus transition zone (2-way ANOVA; side: p<0.0001, F=65.4, DF=1; treatment: p<0.0001, F=57.3, DF=1; interaction: p<0.0001, F=33, DF=1; Bonferroni’s test, carbocaine ipsi 5.1+/-1.5 neurons versus vehicle ipsi 19.3+/-1.1 neurons, p<0.001; Fig.4F).

### 3.3 Effects of Menthol, AITC and cold stimulation on sensory neurons innervating the facial skin and the buccal mucosa

To clarify the participation of TRPM8 and TRPA1 receptor in the detection of cold stimulation, we evaluated neuronal activity in response to ice cooled HBSS (0-2°C), the TRPM8 agonist menthol and the TRPA1 agonist AITC in neurons innervating the oral mucosa or the skin. Because neurons innervating teeth are only a low proportion of the total trigeminal sensory neurons we were not able to detect sufficient numbers of DiI retrolabeled sensory neurons innervating the dental pulp.

A similar proportion of neurons innervating the skin or the mucosa responded to menthol (3.0% and 3.1% respectively) or AITC (56.0% and 46.7% respectively, p=0.10, Chi-square test) and no neurons responded to both menthol and AITC (Fig.5C). A detailed analysis comparing neurons responding to cold stimulation and menthol showed that all neurons responding to menthol were also sensitive to cold stimulation, but a large proportion of neurons were cold sensitive and menthol insensitive (Fig.5D). Moreover, compared to the neurons innervating the skin, a higher proportion of neurons innervating the mucosa were cold sensitive (25.7% and 57.9% respectively, p<0.0001, Chi-square test; Fig.5D). The analysis of neuronal sensitivity to AITC and cold stimulation showed that a subpopulation of cold sensitive neurons was AITC insensitive (10.1% and 37% of neurons innervating the skin or the mucosa respectively; Fig.5E). Conversely, a significant proportion of neurons innervating the skin (15.5%) or the mucosa (21%) were AITC sensitive and cold insensitive, suggesting that TRPA1 is only partially involved in cold perception (Fig.5E).

## References

Bautista DM, Jordt SE, Nikai T, Tsuruda PR, Read AJ, Poblete J, Yamoah EN, Basbaum AI, Julius D. 2006. TRPA1 mediates the inflammatory actions of environmental irritants and proalgesic agents. Cell. 124(6):1269–1282.

Cadden SW, Lisney SJ, Matthews B. 1983. Thresholds to electrical stimulation of nerves in cat canine tooth-pulp with A beta-, A delta-and C-fiber conduction velocities. Brain Res. 261(1):31–41.

Chen J, Joshi SK, DiDomenico S, Perner RJ, Mikusa JP, Gauvin DM, Segreti JA, Han P, Zhang XF, Niforatos W, et al. 2011. Selective blockade of TRPA1 channel attenuates pathological pain without altering noxious cold sensation or body temperature regulation. Pain. 152(5):1165–1172.

Dababneh RH, Khouri AT, Addy M. 1999. Dentine hypersensitivity—an enigma? A review of terminology, epidemiology, mechanisms, aetiology and management. Br Dent J. 187(11):601–611.

Fried K, Arvidsson J, Robertson B, Brodin E, Theodorsson E. 1989. Combined retrograde tracing and enzyme/immunohistochemistry of trigeminal ganglion cell bodies innervating tooth pulps in the rat. Neuroscience. 33:101–9.

Fried K, Sessle BJ, Devor M. 2011. The paradox of pain from tooth pulp: low-threshold “algoneurons”? Pain. 152(12):2685–2689.

Gibbs JL, Melnyk JL, Basbaum AI. 2011. Differential TRPV1 and TRPV2 channel expression in dental pulp. J Dent Res. 90(6):765–770.

Ichikawa H, Deguchi T, Fujiyoshi Y, Nakago T, Jacobowitz DM, Sugimoto T. 1995. Parvalbumin-and calretinin-immunoreactive trigeminal neurons innervating the rat molar tooth pulp. Brain Res. 679(2):205–211.

Itotagawa T, Yamanaka H, Wakisaka S, Sasaki Y, Kato J, Kurisu K, Tsuchitani Y. 1993. Appearance of neuropeptide Y-like immunoreactive cells in the rat trigeminal ganglion following dental injuries. Arch Oral Biol. 38:725–728.

Julius D. 2013. TRP channels and pain. Annu Rev Cell Dev Biol. 29:355–384.

Karashima Y, Talavera K, Everaerts W, Janssens A, Kwan KY, Vennekens R, Nilius B, Voets T. 2009. TRPA1 acts as a cold sensor in vitro and in vivo. Proc Natl Acad Sci U S A. 106(4):1273–1278.

Knowlton WM, Palkar R, Lippoldt EK, McCoy DD, Baluch F, Chen J, McKemy DD. 2013. A sensory-labeled line for cold: TRPM8-expressing sensory neurons define the cellular basis for cold, cold pain, and cooling-mediated analgesia. J Neurosci. 33(7):2837–2848.

Kwan KY, Allchorne AJ, Vollrath MA, Christensen AP, Zhang DS, Woolf CJ, Corey DP. 2006. TRPA1 contributes to cold, mechanical, and chemical nociception but is not essential for hair-cell transduction. Neuron. 50(2):277–289.

McKemy DD, Neuhausser WM, Julius D. 2002. Identification of a cold receptor reveals a general role for TRP channels in thermosensation. Nature. 416(6876):52–58.

McNamara CR, Mandel-Brehm J, Bautista DM, Siemens J, Deranian KL, Zhao M, Hayward NJ, Chong JA, Julius D, Moran MM, et al. 2007. TRPA1 mediates formalin-induced pain. Proc Natl Acad Sci U S A. 104(33):13525–13530.

Nassini R, Gees M, Harrison S, De Siena G, Materazzi S, Moretto N, Failli P, Preti D, Marchetti N, Cavazzini A, et al. 2011. Oxaliplatin elicits mechanical and cold allodynia in rodents via TRPA1 receptor stimulation. Pain. 152(7):1621–1631.

Paik SK, Park KP, Lee SK, Ma SK, Cho YS, Kim YK, Rhyu IJ, Ahn DK, Yoshida A, Bae YC. 2009. Light and electron microscopic analysis of the somata and parent axons innervating the rat upper molar and lower incisor pulp. Neuroscience. 162(4):1279–1286.

Park CK, Kim MS, Fang Z, Li HY, Jung SJ, Choi SY, Lee SJ, Park K, Kim JS, Oh SB. 2006. Functional expression of thermo-transient receptor potential channels in dental primary afferent neurons: implication for tooth pain. J Biol Chem. 281(25):17304–17311.

Patel R, Gonçalves L, Newman R, Jiang FL, Goldby A, Reeve J, Hendrick A, Teall M, Hannah D, Almond S, et al. 2014. Novel TRPM8 antagonist attenuates cold hypersensitivity after peripheral nerve injury in rats. J Pharmacol Exp Ther. 349(1):47–55.

Peier AM, Moqrich A, Hergarden AC, Reeve AJ, Andersson DA, Story GM, Earley TJ, Dragoni I, McIntyre P, Bevan S, et al. 2002. A TRP channel that senses cold stimuli and menthol. Cell. 108(5):705–715.

Story GM, Peier AM, Reeve AJ, Eid SR, Mosbacher J, Hricik TR, Earley TJ, Hergarden AC, Andersson DA, Hwang SW, et al. 2003. ANKTM1, a TRP-like channel expressed in nociceptive neurons, is activated by cold temperatures. Cell. 112(6):819–829.

Yang H, Bernanke JM, Naftel JP. 2006. Immunocytochemical evidence that most sensory neurons of the rat molar pulp express receptors for both glial cell line–derived neurotrophic factor and nerve growth factor. Arch Oral Biol. 51(1):69–78.

